# Intracellular transport of electrotransferred DNA cargo is governed by coexisting ergodic and non ergodic anomalous diffusion

**DOI:** 10.1101/2021.04.12.435513

**Authors:** Aswin Muralidharan, Hans Uitenbroek, Pouyan E. Boukany

## Abstract

The ability of exogenous DNA cargo to overcome the active and viscoelastic eukaryotic cytoplasm is a principal determinant for the gene delivery efficacy. During DNA electrotransfer, DNA forms complexes with the membrane (DNA cargo) which is transported through the cytoplasm through a combination of passive diffusion and active transport. However, this process is poorly understood limiting rational optimization of DNA cargo to be delivered to different cell types. We have investigated the intracellular transport of DNA cargo (of sizes 100 bp, 250 bp and 500 bp) delivered by electrotransfer to non-cancerous and cancerous mammalian cells. We demonstrate that intracellular DNA cargo transport is governed by coexisting ergodic and non ergodic anomalous diffusion for all the tested DNA sizes and cell types. The apparent diffusion coefficient of the electrotransferred DNA cargo in the cytoplasm decreases when the DNA size is increased from 100 bp to 500 bp. Interestingly, the electrotransferred DNA cargo (500 bp) transport is strongly dependent on the cell’s cancer state. Intracellular electrotransferred DNA cargo transport has a higher probability of superdiffusive transport and lower probability of caging in metastatic cells compared to malignant cells followed by benign cells.

## INTRODUCTION

Intracellular transport of exogenous cargo such as nucleic acids through the dense and active intracellular barrier is an indispensable step in biological processes such as cell transfection, up/down-regulation of gene expression, and gene editing [1– 3]. Several physical and chemical strategies are currently employed for cell transfection by delivering nucleic acids such as RNA and DNA [4]. One of the safest and simplest physical methods to introduce the negatively charged nucleic acids to living cells is using pulsed electric fields (also called electrotransfer), which transiently disrupts the cell membrane [3, 4]. During the electrotransfer of nucleic acids to living cells, small nucleic acids like naked siRNA [5] and DNA fragments [6] (15-22 base pair or bp) freely translocate across the permeabilized cell membrane while DNA fragments with size above 25 bp [6] and plasmid DNA (4.7 kbp) [7] form DNA-membrane complexes (or DNA cargo). Direct visualization of electrotransferred plasmid DNA cargo trajectories in the Chinese Hamster Ovary (CHO) cell cytoplasm demonstrated the presence of fast active transport alongside slow passive diffusion in the DNA cargo transport [8]. Classical endocytotic pathways facilitate subsequent transport of the DNA cargo through the cytoplasm [9–11]. Understanding the DNA cargo’s intracellular transport is key to obtaining precise control over DNA electrotransfer efficacy.

What strategies does a cell employ to transport the mesoscopic cargo with a similar size as DNA cargo (0.1 - 0.5 µm [7]) through the cytoplasm? Observations from the intracellular motion of exogenous cargo such as spherical beads [12, 13], carbon nanotubes [14] and nanoparticles [15, 16], and endogenous organelles such as endosomes [17, 18] and insulin granules [19] provide the basis for our current understanding of intracellular mesoscopic cargo transport [20–22]. Geometric constraints such as the cargo size, macromolecule crowding in the cytoplasm, and binding of the cargo to the cytoskeleton hinder diffusion [16, 23–25]. On the other hand, active forces generated by molecular motors on the cargo and cytoplasmic fluctuations can enhance diffusion and often facilitate directed motion [12–14, 18]. Hence, mesoscopic cargo’s intracellular motion often deviates from Brownian motion and is classified as subdiffusive or superdiffusive (or broadly as anomalous diffusion) depending on the randomness of the trajectory [26]. Such classifications, however, are often insufficient to characterize the cargo’s complex motion in the cytoplasm. Mesoscopic cargo such as beads, insulin granules, chromosomal loci, and RNA-protein particles in both eukaryotic and prokaryotic cytoplasm perceive the cytoplasm as a viscoelastic media [12, 16, 27, 28]. Furthermore, the intracellular motion of insulin granules, nanoparticles, and quantum dots in non-cancer and cancer cell lines display ergodicity breaking, or non-equivalence between long-time-averaged motion and ensemble-averaged motion as the cargo can bind to the cytoplasmic components [16, 19, 26, 29]. To rationally optimize the intracellular DNA cargo transport, we should understand its intracellular transport across different DNA sizes (to mimic the geometric constraints) and cell types (with different intracellular activity levels).

In this work, we propose a unified framework for electrotransferred DNA cargo transport in the cytoplasm across different DNA sizes (100 bp, 250 bp, 500 bp) and cell types (animal cell line, benign, malignant, and metastatic breast carcinoma cell line). We demonstrate that the intracellular DNA cargo transport is composed of both ergodic and non-ergodic anomalous diffusion processes (with anomalous exponent α) for all tested DNA sizes and cell types. Increasing the size of the DNA from 100 bp to 500 bp decreases mean apparent intracellular (CHO-K1 cells) diffusion coefficient of the DNA cargo from 1.8 × 10^−2^ µm^2^s^*−α*^ to 1.1 × 10^−2^ µm^2^s^*−α*^. The anomalous exponent of 500 bp DNA cargo motion in cytoplasm is strongly correlated to the cell’s cancer state while the apparent diffusion coefficient is not. Electrotransferred DNA cargo in the cytoplasm has a higher probability to undergo superdiffusive motion in metastatic breast cancer cells compared to malignant followed by benign breast cancer cells.

## RESULTS AND DISCUSSION

### Analysis of intracellular DNA cargo trafficking

To understand the intracellular DNA cargo transport, we study the DNA cargo trajectories after they are internalized (between 20 to 60 minutes after electrotransfer) by the CHO-K1 cells. A representative example of Chinese Hamster Ovary cells (CHO-K1, DSMZ) with internalized fluorescently labeled DNA cargo is shown in Fig. 1*A*. The steric interactions of the DNA cargo with the cytoplasm results in the confinement of the trajectories within the cytoskeletal meshwork, and the biochemical interactions with molecular motors during active transport leads to distinctly longer trajectories (as schematically represented in Fig. 1*B*). As a first step, we used DNA cargo formed by electrotransferred DNA fragments (of sizes 100 bp, 250 bp, and 500 bp) to emulate different levels of steric interactions with the cytoplasm. To quantify the different length scales explored by the DNA cargo, we estimate the radius of gyration **R**_**g**_(*T*) of the individual trajectories, where 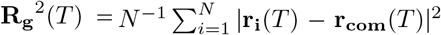 where T is the measurement time [30]. The individual trajectories were sampled as *N* points with positions **r**_**i**_ and has a center of mass **r**_**com**_ (assuming same mass at all points). The probability densities of |**R**_**g**_| for *T* = 20 s of the DNA cargo trajectories for the different DNA fragment sizes is plotted in Fig. 1*C* -*E*. Majority of the DNA cargo display localized displacements (|**R**_**g**_| ∼ 75 nm) for all DNA sizes. The peak of the probability density distribution from |**R**_**g**_| of the trajectories is close to the characteristic actin mesh size (100-200 nm) in a living cell [31]. The DNA cargo traverse up to |**R**_**g**_| ∼ 950 nm within 20 seconds for all DNA fragment sizes. The localized and exploratory intracellular electrotransferred DNA cargo transport was previously reported in plasmid DNA cargo delivered to CHO cells and is dependent on the cell’s cytoskeleton state [8].

**FIG. 1.**
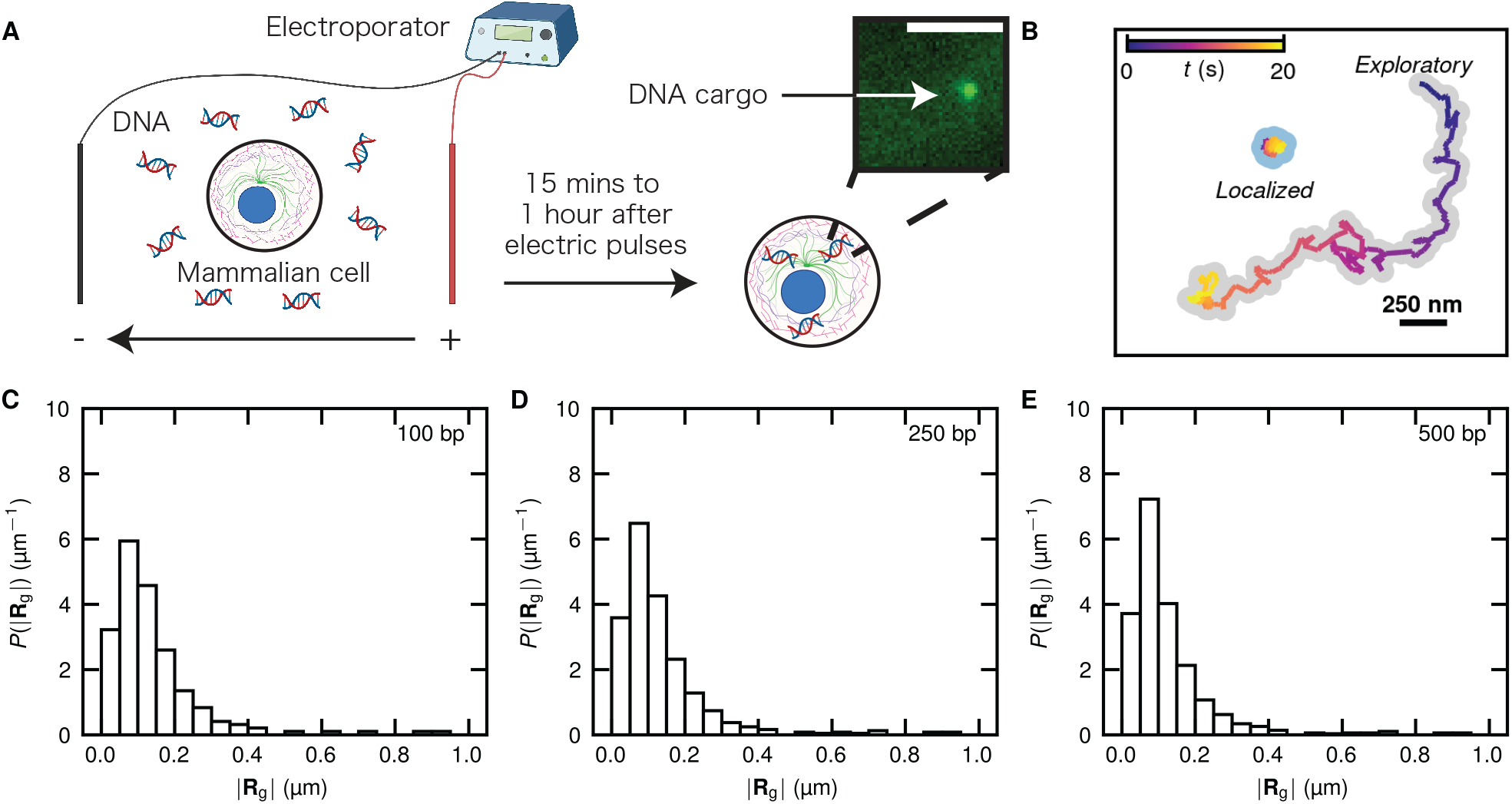
Different intracellular DNA cargo transport modes. (*A*) A schematic of delivery of DNA cargo to mammalian cells. Ten electric pulses of 350V/cm with a duration of 5 ms and a frequency of 1 Hz is applied to deliver YOYO-1 labelled DNA fragment to mammalian cells. DNA-membrane complex (DNA cargo) is formed and is internalized by the cell. The direct visualization of the intracellular DNA cargo transport is performed by fluorescent microscopy between 15 minutes and 1 hour after the electric pulses are applied. A representative fluorescent image of 100 bp DNA cargo inside a CHO-K1 cell is shown. The scale bar represents 5 µm. The schematic is created with BioRender. (*B*) Two example trajectories displaying localized motion covering small distances and exploratory motion covering larger distances. The trajectories are colored according to the time from the starting position. (*C*)-(*E*) The probability density distribution (bin size = 0.05 µm) of the radius of gyration **R**_**g**_ (*T* = 20 s) of the DNA cargo trajectories for (*C*) 100 bp (*n* = 191 trajectories from 38 cells), (*D*) 250 bp (*n* = 99 trajectories from 39 cells), (*E*) 500 bp (*n* = 210 trajectories from 34 cells) DNA in CHO-K1 cell cytoplasm.

Having established that the DNA cargo can explore different length scales, we investigate its one dimensional displacement at different lag times. For a particle performing ideal random Brownian walk in a simple Newtonian fluid, the probability density of one dimensional displacement *P* (|Δ*x*|, Δ*t*) exhibits a Gaussian distribution (*P* (|Δ*x* |) ∼ exp(−|Δ*x*| ^2^ */σ*), where *σ* is a scale parameter) by the virtue of the central limit theorem [32]. One-dimensional displacement Δ*x* is defined as *x*(*t* + Δ*t*) − *x*(*t*), where *x*(*t*) is the DNA position at time *t* (in either *x* or *y* direction), and Δ*t* is the time interval between the DNA position measurements. The displacements in *x* and *y* directions are combined to improve statistics. We plot the probability density of the absolute value of the intracellular one dimensional DNA cargo displacements, *P* (|Δ*x*|, Δ*t*), for Δ*t* = 0.1 s is plotted in Figs. 2*A*-*C*. *P* (|Δ*x*|, Δ*t*) follows a Laplacian distribution (*P* (|Δ*x*|) ∼exp(−|Δ*x*| */σ*)) for the majority of the distribution, displaying enhanced probability for the cargo to make large displacements compared to a regular Brownian motion. The cytoplasmic heterogeneity (which results in a broad distribution of diffusivities of particles within the cytoplasm), non-thermal active fluctuations and fast active endocytotic transport of DNA cargo [9] results in a deviation from Gaussianity due to the co-existence of fast and slow particles [28]. We quantify the deviation from Gaussian behavior by excess kurtosis, *κ* = ⟨Δ*x*^4^⟩*/* ⟨Δ*x*^2^⟩^2^ −3. For a Gaussian distribution, *κ* = 0, and distribution with longer tails compared to Gaussian display *κ >* 0. *P* (|Δ*x* |, Δ*t*) remains leptokurtic (*κ >* 0) for all DNA sizes. If the long exponential tails in the displacement probability distribution arise from continuous drift (∼Δ*t*) from active transport, *κ* would grow with lag time as the exponential tails would grow faster compared to its passive diffusion counterpart represented by a Gaussian 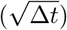. Fig. 2*D* shows that non-Gaussian statistics is maintained even at longer lag times. 100 bp DNA cargo showed an increase in *κ* with Δ, while 250 bp DNA cargo showed no visible correlation with Δ*t* and 500 bp DNA cargo showed a decrease in *κ* with Δ*t*. We find that the larger displacements appear as intermittent spikes rather than continuous drift with the same velocity from the DNA cargo trajectories (Fig. S1, supplementary information). Such bursts of large active transport are consistent with the view that endosomal transport involves stress buildup in the cytoskeleton followed by a rapid release and can explain the weak correlation of *κ* with Δ*t* [17]. We conclude that the physical mechanism that can describe the intracellular DNA cargo transport should have non-Gaussian displacements. We investigate the influence of the DNA size on the DNA cargo’s cytoplasmic mobility in the following section.

**FIG. 2.**
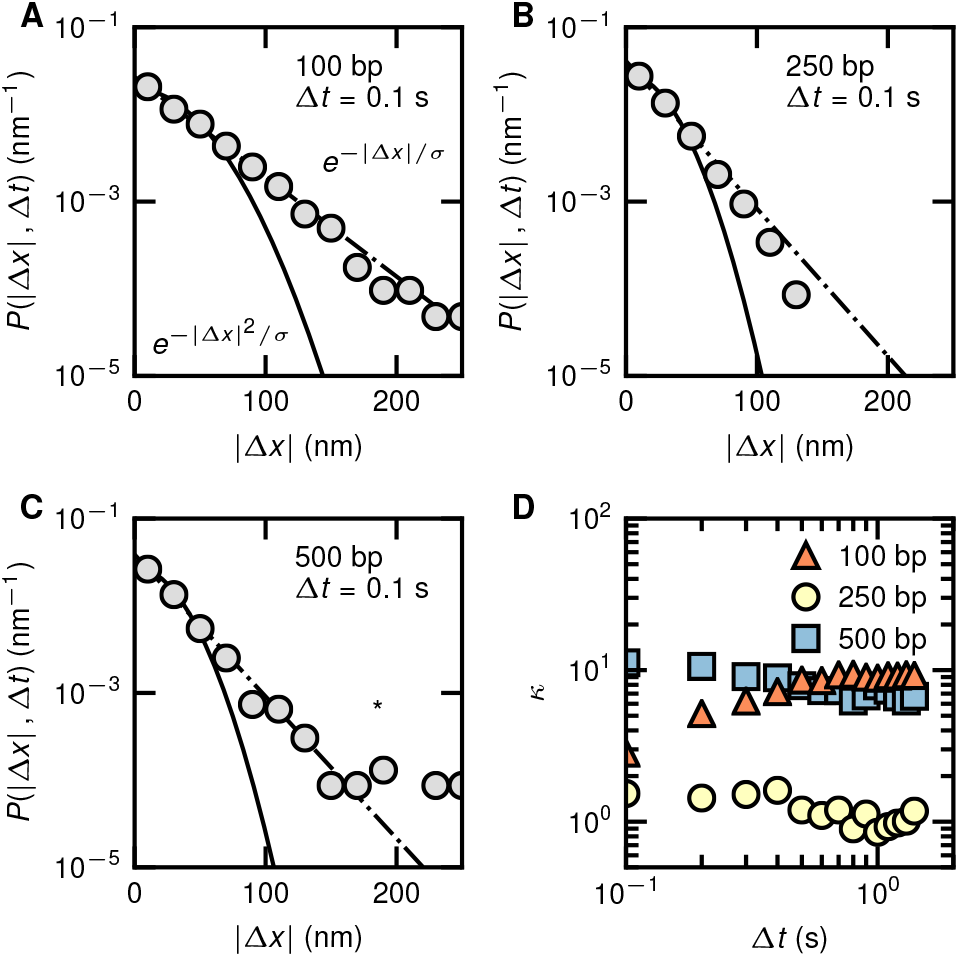
Non-Gaussian DNA cargo displacements. Probability density distribution (bin size = 20 nm) for one dimensional intracellular DNA cargo displacement during a time interval of Δ*t* = 0.1 s is plotted for (*A*) 100 bp (*B*) 250 bp and (*C*) 500 bp DNA. The displacements in *x* and *y* directions are collected together for better statistics. The solid line represents a Gaussian fit to the center of the distribution while the dash-dot line represents a Laplacian fit to the entire distribution. The asterisk (*) in (*C*) represents the location where the distribution deviates from the Laplacian distribution. (*D*) Deviation from Gaussianity is represented by excess kurtosis *κ* for different Δ*t* for the 100, 250 and 500 bp DNA.

### Intracellular anomalous diffusion of DNA cargo

To evaluate how the cytoplasmic mobility of DNA cargo varies with DNA size, we estimate the relation between mean square displacement

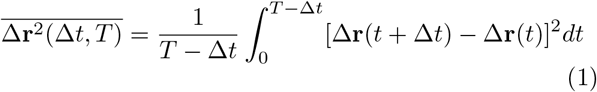

and lag time Δ*t* for a measurement time *T* = 10 s. 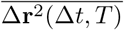 is plotted against Δ*t* for *T* = 10 s in Figs. 3*A*-*C* to characterize the intracellular transport. Individual 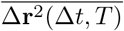 trajectories for all the sizes of DNA (colored in gray) is spread over two orders of magnitude demonstrating that the cargo trajectories are heterogeneous. 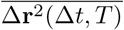 increases with Δ*t* and deviate from linearity. Furthermore, 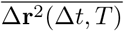 display two distinct scaling with Δ*t*, with 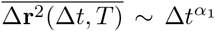 when Δ*t* < 1 s, and 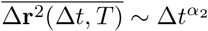 when Δ*t >* 1 s where *α*_1_ and *α*_2_ are power law exponents and Δ*t* ∼1 s is a characteristic cross-over lag time. Mean square displacement from individual trajectories are fitted to 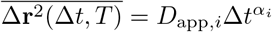 (*i* = 1, 2) to determine the apparent diffusion coefficient *D*_app_ and the anomalous power-law exponent using non-linear regression. *α*_1_ is determined during the interval Δ*t* = 0.1 s and Δ*t* = 0.8 s while *α*_2_ is determined during the interval Δ*t* = 2 s and Δ*t* = 5 s to avoid biases at the edge of the cross-over lag time. Fig. 3*D*-*E* shows that the trajectories display marked heterogeneity in both *D*_app,*i*_ and *α*_*i*_ estimated before and after the cross-over lag time. *D*_app,*i*_ is spread over two to three orders of magnitude as seen in Fig. 3*D* (the whole population is presented as in Fig. S4, supplementary information as a 2D density plot) for all DNA sizes. Fig. 3*D* shows that there is no statistically significant (*P >* 0.05) difference between *D*_app,1_ and *D*_app,2_ within each DNA size. The arithmetic mean of the diffusion coefficient ⟨*D*_app,1_⟩decreases from ∼1.8 × 10^−2^ µm^2^s^*−α*^ (standard error of the mean, s.e.m. ∼2 × 10^−3^ µm^2^s^*−α*^) for 100 bp DNA to ∼1.1 × 10^−2^ µm^2^s^*−α*^ (s.e.m. ∼1.4 × 10^−3^ µm^2^s^*−α*^) for 500 bp DNA (*P* < 0.01). Similarly, the arithmetic mean of the diffusion coefficient ⟨*D*_app,2_⟩decreases from ∼2 × 10^−2^ µm^2^s^*−α*^ (standard error of the mean, s.e.m. ∼3.8 × 10^−3^ µm^2^s^*−α*^) for 100 bp DNA to ∼1.2 × 10^−2^ µm^2^s^*−α*^ (s.e.m. ∼2.7 × 10^−3^ µm^2^s^*−α*^) for 500 bp DNA (*P* < 0.01). No significant differences in *D*_app,*i*_ were observed between 100 bp and 250 bp DNA cargo, and 250 bp and 500 bp DNA cargo. Our results suggest that viscous dissipation due to increasing DNA cargo size might be responsible for reduction in the diffusion coefficient.

**FIG. 3.**
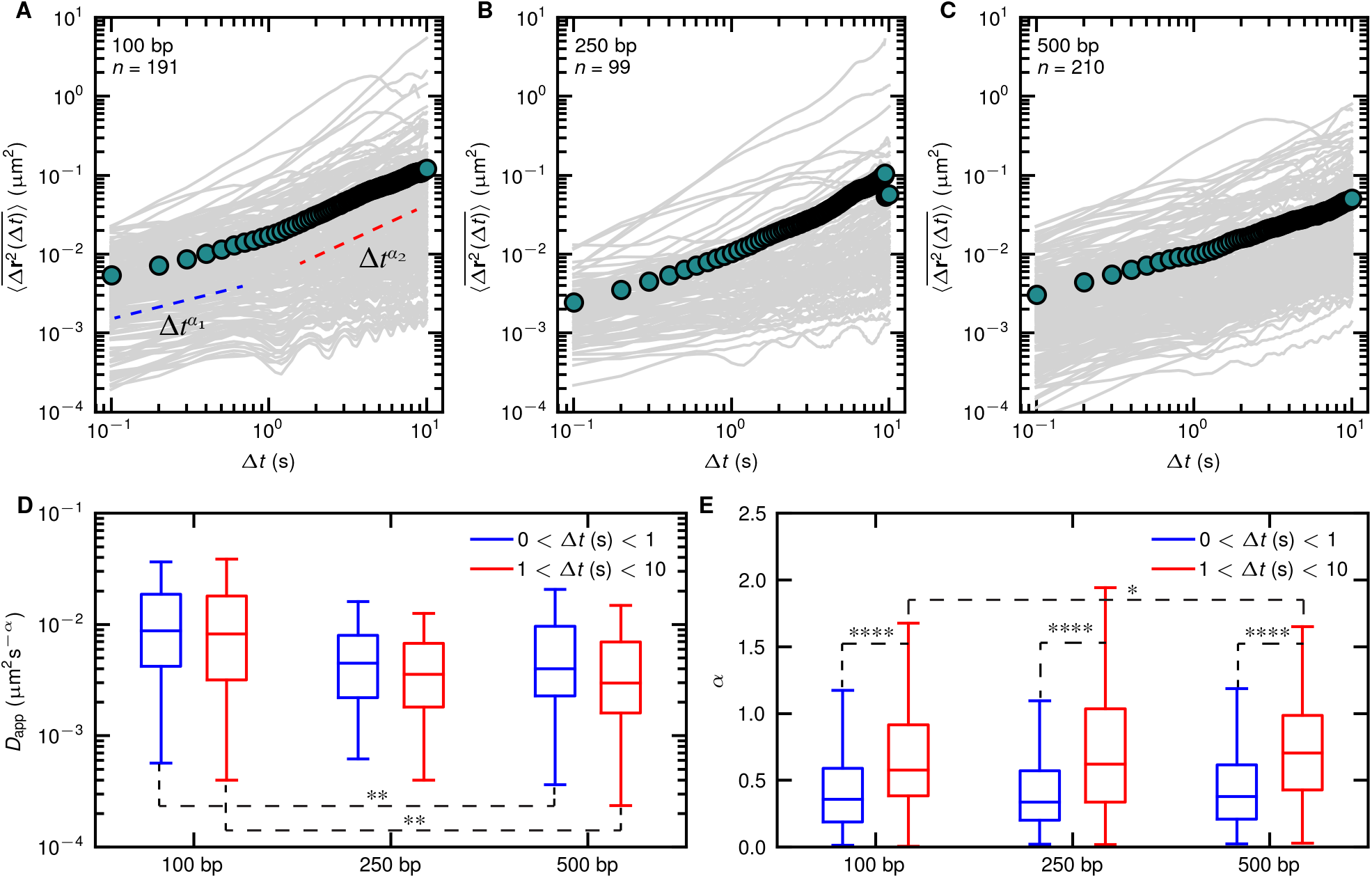
Intracellular anomalous DNA cargo transport. Individual time averaged mean square displacements 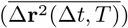 are plotted in solid gray color for (*A*) 100 bp (*B*) 250 bp (*C*) 500 bp DNA against the lag time Δ*t*, for a measurement time *T* = 10 s. The green circles represent the ensemble and time average mean square displacement 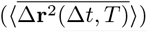. (*D*) The apparent diffusion coefficient *D*_app_, and (*E*) anomalous exponent *α* obtained by fitting the individual 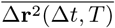 to 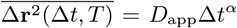 before and after the cross-over Δ*t* = 1 s is plotted as a box plot for different DNA sizes. Statistical significance of difference of the mean was tested using one way analysis of variance. *: *P* < 0.05, **:*P* < 0.01, ***: *P* < 0.001, ****: *P* < 0.0001. Non significant difference between the datasets were not displayed for better clarity of the figure.

To classify the randomness of the trajectory, we then analyse the anomalous exponents *α*_*i*_ for the different DNA fragment sizes. The value of *α*_*i*_ contains information about the driving forces of the cargo transport. For Brownian motion *α*_*i*_ = 1 [33]. The motion of the cargo in a viscoelastic or crowded environment like the cytoplasm result in *α*_*i*_ < 1. Only motion with non-thermal driving force, for e.g., active forces can lead to superdiffusive motion with an anomalous exponent *α*_*i*_ *>* 1. Both *α*_1_ and *α*_2_ display trajectory to trajectory fluctuations for all sizes of DNA, and showed either caged (*α*_*i*_ < 0.4), subdiffusive (0.4 < *α*_*i*_ < 1) and superdiffusive power law scaling as seen in Fig. 3*E*. Approximately 60 % of the trajectories are caged, ∼35 % subdiffusive when Δ*t* < 1 s as shown in Fig. S5 (supplementary information) for all sizes of DNA. For all the DNA sizes, the anomalous exponent showed a significant (*P* < 0.0001) increase after the cross-over lag time of 1 s. The anomalous exponent before Δ*t* ∼1 s was independent of DNA size. After the cross-over Δ*t* ∼1 s, majority of the trajectories (∼60%) are subdiffusive, ∼20% caged and ∼20% superdiffusive. After Δ*t* ∼1 s, there was a significant (*P* < 0.05) increase in the anomalous exponent between 100 bp DNA cargo and 250 bp DNA cargo. No significant differences were observed between other combinations of DNA size. The arithmetic mean of the anomalous exponents ⟨*α*_1_⟩ ∼0.42 (s.e.m. < 0.03) for all the DNA sizes and ⟨*α*_2_⟩ ∼0.61 (s.e.m. ∼0.03) for 100 bp, ∼0.64 (s.e.m. ∼0.05) for 250 bp and ∼0.72 (s.e.m. ∼0.03) for 500 bp DNA. Fig. 3*E* shows that the median *α* increases from ∼0.4 to ∼0.7 after the cross-over Δ*t* ∼1 s. The ensemble and time averaged mean square displacement 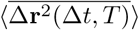 also show an increase in anomalous exponent after Δ*t* ∼1 s for all DNA sizes.

What mechanisms lead to the observed anomalous diffusion of the DNA cargo? Multiple molecular motors drag the DNA cargo encapsulated in endosomes [9] along microtubule tracks through viscoelastic cytoplasm [18]. The intracellular motion of endosomes therefore involve stress buildup in the cytoplasm followed by bursts of active motion [17]. The increasing size of the cargo is reported to increase the resistance faced from the cytoplasm and can stall the motion while the cooperative action of molecular motors facilitates directionality [18]. Increasing DNA size from 100 bp to 500 bp showed a decrease (∼64%) in the apparent diffusion coefficient *D*_app,1_. A plausible explanation for this observed decrease in *D*_app,*i*_ is the potential increase in the cytoplasmic resistance due to the larger cargo size. In contrast, despite the expectation that the larger cargo enhances the cytoplasm’s resistance and decreases the probability for exhibiting directionality and superdiffusive motion, increasing the DNA size from 100 bp to 500 bp increased *α*_2_. Other combinations of DNA sizes had no statistically significant differences in *α*_*i*_. To further investigate the influence of the DNA size on the randomness of the DNA cargo trajectory, we estimate the directional change angle *θ*(*t*, Δ*t*) defined such that

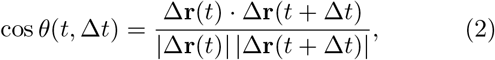

where the lag time Δ*t* represents the temporal coarsening and Δ**r** is the displacement vector [34]. The directional change probability density distribution is plotted in Fig. S3 (supplementary information). At short temporal coarsening (Δ*t* = 0.1 s), the distribution shows a peak at *π* rad, indicating that an anti-persistent motion is dominant at short time scales and lead to the observed caged like motion before the cross over lag time. At further increase in the degree of temporal coarsening, the trajectories become more directional, increasing the anomalous exponent after the cross over lag time(Fig. S3*D*). However, the directional change probability density distribution and DNA size had no statistically significant correlation, which leads us to conclude that the DNA size does not significantly alter molecular motor’s ability to overcome the cytoplasm’s viscoelastic resistance and actively transport the electrotransferred DNA cargo. We probe into the physical models of anomalous diffusion which can best describe DNA cargo transport in the following section. To do so, we check for ergodicity breaking in the DNA cargo dynamics.

### Coexistence of non-ergodic and ergodic processes in the intracellular DNA cargo dynamics

To understand if the statistical properties obtained from individual trajectories during the observation time represent the ensemble behavior, we test if the DNA cargo motion is ergodic. As a first step towards checking for ergodicity, we plot the time-averaged mean squared displacement for individual trajectories for different measurement times *T* and lag time Δ*t* = 0.1 s in Figs. 4*A*-*C* [35]. We use overlines to denote time averages and angular brackets to denote ensemble averages. The mean square displacements of different cargo trajectories at the 0.1 s interval are scattered over three orders of magnitude for all tested DNA sizes. The scatter of mean square displacement is more pronounced at short measurement times due to the lower denoising strength from lower degree of averaging. As the measurement time increases, the number of points per sample in averaging increases reducing the noise in the mean square displacement. The dependence of 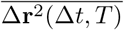 on the measurement time *T* is analyzed by fitting the data to a power law of the form *T* ^*β*^ for each trajectory, where *β* is the power-law exponent. The fitting is performed between *T* = 0.8 s and *T* = 10 s to avoid errors due to noise in the data at low measurement times. The probability distribution of *β* for the different sizes of DNA is provided in Fig. S2 (supplementary information). For most of the trajectories for all DNA sizes, *P* (*β*) ∼ *T*^0^. Some of the trajectories display a decrease while some display an increase in 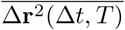 with *T*. Despite the strong heterogeneity at the single trajectory level, the ensemble average 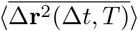 is independent of the measurement time (∼*T*^0^). The power-law dependence of the mean square displacement with measurement time is often seen in non-ergodic systems. To quantify the degree of amplitude fluctuations of individual 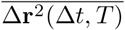, the ergodicity breaking parameter

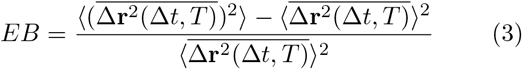

is calculated [35–37]. For an ergodic process, *EB* = 0 when *T* → ∞ as there is no amplitude fluctuations in 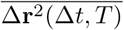 [26]. Fig. 4*D* shows that *EB* converges to *EB* ∼1 with increasing measurement time for 100 bp and 250 bp DNA and *EB* ∼2 for 500 bp DNA, showing that intracellular DNA motion after electroporation is non-ergodic.

**FIG. 4.**
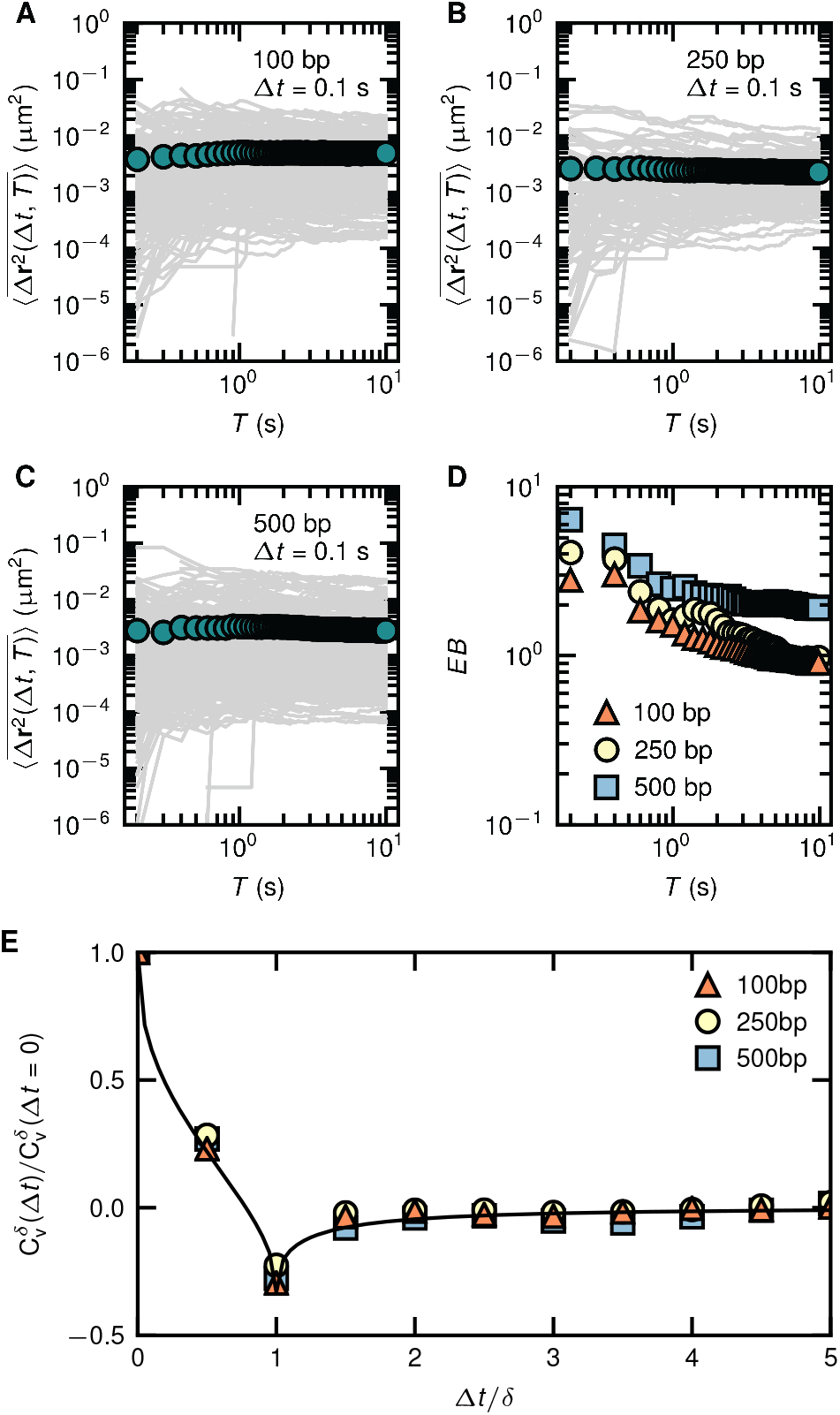
Coexistence of ergodic and non-ergodic processes in the intracellular DNA cargo dynamics. (*A*)-(*C*) Individual time averaged mean square displacements 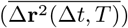 are plotted in solid gray color for (*A*) 100 bp (*B*) 250 bp (*C*) 500 bp DNA against the measurement time *T*, for a lag time Δ*t* = 0.1 s. The green circles represent the ensemble and time average mean square displacement 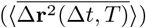. (*D*) The ergodicity breaking parameter *EB* is plotted against the measurement time *T*. (*E*) Velocity auto-correlation function 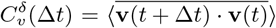, where **v**(*t*) = [**r**(*t* + *δ*) − **r**(*t*)]*/δ* normalized by 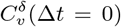 plotted against the lag time Δ*t* normalized by the discretization time interval *δ* = 0.2 s for 100, 250 and 500 bp DNA. The solid line is the analytical prediction from fractional Brownian motion (FBM), Eq. 5 with ⟨*α*⟩ = 0.42.

Can a unified framework describe the DNA cargo’s non ergodic anomalous diffusion with non Gaussian displacements? Simple Brownian walk slowed down by molecular crowding in the cytoplasm is insufficient to explain the observed heterogeneity in the mean square displacement. Continuous time random walk [26] (CTRW), where the trajectories consist of random steps which are broadly distributed in time or fractional Brownian motion [38] (FBM), where the trajectories consist of random steps which are broadly distributed in space are often used to model heterogenous intracellular mesoscopic cargo motion. A distinguishing statistical property of CTRW from FBM is that CTRW is non-ergodic. During biological processes like intracellular insulin granule transport in pancreatic β-cells [19] and membrane diffusion of voltage gated potassium channels in human embryonic kidney cells [39], ergodic FBM and non-ergodic CTRW coexist. While the ergodicity breaking suggests that DNA cargo motion follow a CTRW, the DNA cargo mean square displacement does not plateau as typically observed in a CTRW. Furthermore, we did not find any evidence for intermittent stalling in our trajectories (Fig. S1, supplementary information) which is a signature of CTRW. These observations lead us to hypothesise that an ergodic FBM process coexists with the CTRW. To check whether our data can be described by ergodic FBM, we proceed to calculate the velocity auto-correlation function

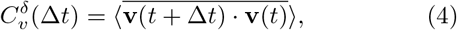

where **v**(*t*) = [**r**(*t* + *δ*) − **r**(*t*)]*/δ*, for discretization time interval *δ* = 0.2 s [27, 40, 41]. Fig. 4*E* shows that all the DNA sizes have a negative autocorrelation 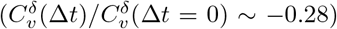 or memory at Δ*t/δ* = 1 before levelling off at zero. The velocity autocorrelation is independent of the DNA size used. Such anticorrelated motion is exhibited by cargo undergoing ergodic fractional Brownian walk (FBM) [42]. Fig. 4*E* shows that the velocity anticorrelation function shows good agreement (root mean square error = 0.29) with the analytical predictions of FBM [25, 26]

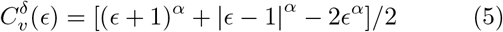

where *ϵ* = Δ*t/δ*, with *α* set to the value ⟨*α*_1_⟩ = 0.42 (obtained in the previous section). The velocity autocorrelation function is self similar over discretization time interval as shown in Fig. S6 (supplementary information). The directional change distribution plotted in Fig. S3*A* (supplementary information) which shows a peak directional change of *π* rad at Δ*t* = 0.1 s corroborates the short lag time (∼ Δ*t* = 0.1 s) anti-persistent motion. At further temporal coarsening (Δ*t* = 0.5 s and Δ*t* = 1 s), the peak shifts towards 0 rad and 2*π* rad indicating emergence of directional inertial motion at long lag times (Fig. S3*B* -*D*, supplementary information).

Overall, our data presents evidence that ergodic and non-ergodic processes coexist in intracellular electrotransferred DNA cargo transport. Non-ergodic processes like CTRW originate from transient binding and unbinding of the molecular motors which are attached to the cargo with different components of the cytoplasm. Our experiments display characteristic signatures of CTRW including ergodicity breaking and non-Gaussianity. The ergodic FBM characteristics exhibited by the DNA cargo potentially originates from the intracellular fluctuations. Hence, both the cargo and the cytoplasmic environment potentially govern the intracellular DNA cargo dynamics. To keep the permeabilization area during electroporation constant in our studies, we did not chemically perturb the cytoskeleton and the molecular motors [43]. In the following section, we study the influence of different levels of cytoplasmic activity (by using breast cancer cells at different stages of cancer) on the intracellular DNA cargo transport.

### Influence of cell’s state on the electrotransferred DNA cargo transport

To understand the influence of the cell’s cancer state on the DNA cargo transport, we studied the intracellular transport of 500 bp DNA cargo through breast cancer cells of varying activity. We chose the following cell lines, which have an increasing level of intracellular activity: MCF10A (benign), MCF7 (malignant), and MDA-MB-231 (metastatic) [13, 44]. Ergodicity breaking in the DNA cargo transport is present in all the cancer cell lines with varying activity levels in Fig. 5*A*. Like CHO-K1 cells, *EB* for 500 bp DNA cargo trajectories of all the cancer cell lines studied converges to *EB* ∼ 2 −3. The FBM analytical solution describes the velocity autocorrelation function well for the DNA cargo transport in the cancer cells as shown in Fig. 5*B*. Our combined results from animal and human breast cancer (at different cancer states) cell lines lead us to propose that coexisting ergodic, and non-ergodic transport could be a universal strategy employed by eukaryotic adherent cell lines for intracellular transport of electrotransferred DNA cargo.

**FIG. 5.**
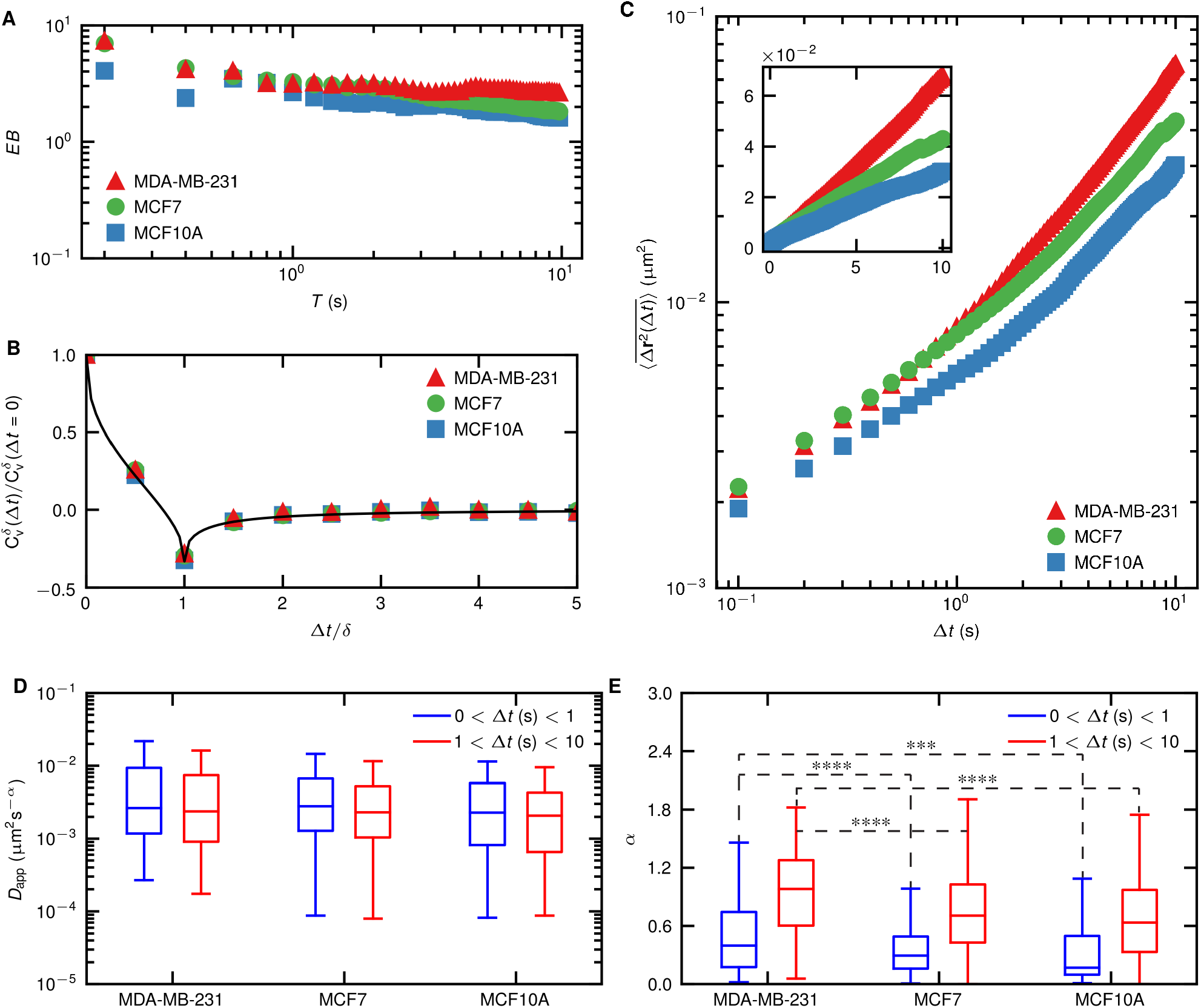
Influence of the cell’s cancer state on the intracellular 500 bp DNA cargo dynamics. MCF10A cells are benign, MCF7 are malignant and MDA-MB-231 cells are metastatic breast cancer cells. (*A*) Ergodicity breaking parameter *EB* estimated for Δ*t* = 0.1 s is plotted against measurement time *T*. (*B*) Velocity auto-correlation function 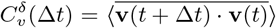, where **v**(*t*) = [**r**(*t* + *δ*) **r**(*t*)]*/δ* normalized by 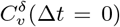 plotted against the lag time Δ*t* normalized by the discretization time interval *δ* = 0.2 s for the cancer cell lines. The solid line is the analytical prediction from fractional Brownian motion (FBM), Eq. 5. *C* Ensemble and time averaged mean square displacement 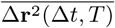 is plotted against the lag time Δ*t* for the cancer cell lines (measurement time *T* = 10 s). The plot is shown in linear scale in the inset. (*D*) The apparent diffusion coefficient *D*_app_, and (*E*) anomalous exponent *α* obtained by fitting the individual 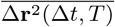 to 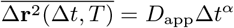 before and after the cross-over Δ*t* = 1 s is plotted as a box plot for different cancer cell lines. Statistical significance of difference of the mean was tested using one way analysis of variance. *: *P* < 0.05, **:*P* < 0.01, ***: *P* < 0.001, ****: *P* < 0.0001. *α* between the two time scales are significantly different for all cell lines (*P* < 0.0001) but are not shown for better clarity. Non significant difference between the datasets were not displayed for better clarity of the figure. 133 trajectories (from 63 cells) were analyzed for MDA-MB-231 cells, 628 trajectories (from 98 cells) were analyzed for MCF7 cells, and 130 trajectories (from 60 cells) were analyzed for MCF10A cells.

To study the influence of the cell’s intrinsic activity on the cytoplasmic mobility of the DNA cargo we plot the 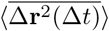 for with measurement time *T* = 10 s in Fig. 5*C*. 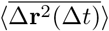 increases from ∼ 3 × 10^−2^ µm^2^ in benign cells to ∼4 × 10^−2^ µm^2^ in malignant cells and ∼ 7 × 10^−2^ µm^2^ in metastatic cells at Δ*t* = 10 s. This shows that electrotransferred DNA cargo traverse longer distances in the cytoplasm of the cells with higher activity. The individual time averaged mean squared displacement show strong heterogeneity over different cell lines as shown in Fig. S7 (supplementary information). The two distinct scaling with Δ*t* with a cross-over Δ*t* ∼ 1 s is present for the 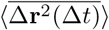 of DNA cargo in cancer cell lines similar to the experiments performed with CHO-K1 cells. *D*_app,*i*_ and *α*_*i*_ govern the relation of 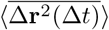 with Δ*t*. To understand why cells with higher intracellular activity have higher 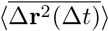, we estimate *D*_app,*i*_ and *α*_*i*_ for the cancer cell lines. The ensemble average ⟨*D*_app,1_⟩ for the 500 bp DNA cargo is ∼8.1 × 10^−3^ µm^2^s^*−α*^ (s.e.m ∼ 1.2 × 10^−3^ µm^2^s^*−α*^) for MDA-MB-231 cells, ∼7.4 × 10^−3^ µm^2^s^*−α*^ (s.e.m ∼ 0.7 × 10^−3^ µm^2^s^*−α*^) for MCF7 cells, and ∼5.7 × 10^−3^ µm^2^s^*−α*^ (s.e.m ∼ 0.8 × 10^−3^ µm^2^s^*−α*^) for MCF10A cells. The ensemble average ⟨*D*_app,2_⟩ for the 500 bp DNA cargo is ∼8.9 × 10^−3^ µm^2^s^*−α*^ (s.e.m ∼1.5 × 10^−3^ µm^2^s^*−α*^) for MDA-MB-231 cells, ∼7.6 × 10^−3^ µm^2^s^*−α*^ (s.e.m ∼0.7 × 10^−3^ µm^2^s^*−α*^) for MCF7 cells, and ∼5.3 ×10^−3^ µm^2^s^*−α*^ (s.e.m ∼0.7 × 10−3 µm^2^s^*−α*^) for MCF10A cells. There is no significant difference (*P >* 0.05) in *D*_app,*i*_ between the different cancer cell lines as shown in Fig. 5*D*. On the other hand, *α*_*i*_ is dependent on the cell’s intrinsic activity as shown in Fig. 5*E*. Both *α*_1_ (*α* when 0 < Δ*t* (s)< 1) and *α*_2_ (*α* when 1 < Δ*t* (s)< 10) is the highest for MDA-MB-231 cells, followed by MCF7 cells and MCF10A cells respectively. The ensemble average ⟨*α*_1_⟩ is 0.49 (s.e.m. ∼0.03) for MDA-MB-231 cells, 0.36 (s.e.m. ∼0.01) for MCF7 cells and 0.33 (s.e.m. ∼0.03) for MCF10A cells. The ensemble average ⟨*α*_2_⟩ is 0.88 (s.e.m. ∼0.04) for MDA-MB-231 cells, 0.7 (s.e.m. ∼0.02) for MCF7 cells and 0.67 (s.e.m. ∼0.04) for MCF10A cells. These observations provide direct evidence that intracellular DNA cargo experience are less subdiffusive in metastatic cell cytoplasm followed by malignant and benign cells respectively. The DNA cargo from all the cell lines experience a short time scale caging (before Δ*t* ∼ 1 s) followed by escape from the cage (Δ*t* ∼ 1 s). The percentage of trajectories which follow caged, subdiffusive and superdiffusive transport are plotted in Fig. S10 (supplementary information). Before Δ*t* ∼1 s, ∼50% of the DNA cargo trajectories are caged, ∼40% are subdiffusive and ∼10%are superdiffusive for MDA-MB-231 cells. After Δ*t* ∼1 s, ∼15% of the DNA cargo trajectories are caged, ∼35%are subdiffusive and ∼50% are superdiffusive for MDA-MB-231 cells. Similarly for MCF7 and MCF10A cells, ∼60% of DNA cargo trajectories are caged, ∼35% are subdiffusive and ∼5% are superdiffusive before Δ*t* ∼1 s. For MCF7 cells, ∼20% of the DNA cargo trajectories are caged, ∼55% are subdiffusive and ∼25% are superdiffusive after Δ*t* ∼1 s. For MCF10A cells, ∼30% of the DNA cargo trajectories are caged, ∼50% are subdiffusive and ∼20% are superdiffusive after Δ*t* ∼1 s. Hence, the percentage of the DNA cargo trajectories that display superdiffusive behavior is strongly correlated to the cancer state of the cell.

What leads to the differences in the anomalous diffusion between metastatic cells, malignant and benign cells? Metastatic cells have a higher degree of intracellular activity followed by malignant cells and benign cells [13, 44]. Since the size of the DNA cargo is the same (as demonstrated by lack of significant differences in *D*_app_ of 500 bp DNA cargo in different cell lines), the observed behavior is strongly correlated to the forces exerted by the molecular motors [18]. Although the cytoplasm remains viscoelastic (as demonstrated by the velocity auto-correlation function) for the DNA cargo for all the cancer cells tested, the values of *α*_1_ suggest that the intracellular fluctuations are significantly higher and leads to the lower degree of caging observed in metastatic cells followed by malignant and benign cells. The long time scale behavior (after Δ*t* ∼1 s) suggests that the forces exerted by the molecular motors on the DNA cargo is higher in the metastatic cells followed by malignant cells and benign cells and leads to a higher fraction of DNA cargo trajectories displaying superdiffusive behavior. The ergodicity breaking of the DNA cargo motion supports the molecular motor’s involvement in the intracellular DNA cargo trajectories. Our results show that the intracellular DNA cargo trajectories in cancerous and non-cancerous cell lines can be described by coexisting ergodic and non-ergodic anomalous diffusion. The intracellular mobility differences across various cell types for a fixed size DNA cargo arise from the cell’s intracellular activity. Overall, our results strongly suggest that the transport of DNA cargo is governed active stresses in the cell cytoplasm (which is governed by the cell’s cancer state) and viscous dissipation in the cytoplasm (which is governed by the DNA size).

## CONCLUSIONS

We demonstrate that the electrotransferred DNA cargo undergoes coexisting ergodic and non-ergodic anomalous diffusion for different DNA sizes (100 bp, 250 bp, 500 bp) and non-cancerous and cancerous cell lines (CHO-K1, MCF10A, MCF7, and MDA-MB-231). The DNA cargo trajectories can be described by anomalous diffusion consisting of two characteristic scalings with lag time and has a crossover lag time ∼1 s. The apparent diffusion coefficient of the DNA cargo in CHO cell cytoplasm decreased when we increased the DNA size from 100 bp to 500 bp. At short lagtimes, most of the DNA cargo trajectories are caged and subdiffusive, while at long lag times, the DNA cargo trajectories are caged, subdiffusive, or superdiffusive. The fraction of DNA cargo trajectories displaying caged, subdiffusive, or superdiffusive behavior has no visible correlation with the DNA size in CHO cell cytoplasm but strongly correlates with the cell’s cancer state. 500 bp DNA cargo in metastatic cytoplasm experience a higher percentage of superdiffusive transport and lowest degree of caging in comparison to malignant cells and benign cancer cell cytoplasm. Hence the viscous dissipation and active cytoplasmic forces are important factors in electrotransferred DNA cargo’s intracellular motion. Our findings provide guidelines to develop new theoretical models to rationally design and optimize DNA electrotransfer protocols for different DNA sizes and cell types.

## I. MATERIALS AND METHODS

### Cell culture

The Chinese Hamster Ovary cells, CHO-K1 (DSMZ), were grown in T-flasks containing culture medium consisting of Nutrient Mixture Ham’s F-12 (Sigma Aldrich) supplemented with ∼10% fetal bovine serum (Sigma Aldrich). MCF-7 (DSMZ) cells were cultured in Dulbecco’s minimal essential medium (Sigma Aldrich) supplemented with ∼10% fetal bovine serum and 0.01 mg/ml human recombinant insulin (Gibco). MCF10A (ATCC) cells were cultured in DMEM/F12 Ham’s Mixture (Gibco) supplemented with ∼5% horse serum (Gibco), 20 ng/ml human epidermal growth factor (EGF) recombinant protein (Invitrogen), 10 µg/ml human recombinant insulin (Gibco), 0.5 µg/ml hydrocortisone (Sigma Aldrich), 0.1 µg/ ml cholera toxin (Sigma Aldrich). MDA-MB-231 (ATCC) cells were cultured in DMEM supplemented with ∼10% fetal bovine serum.

The cells were incubated at 37 ^°^C with 5% CO_2_ and were sub-cultured every two days. Twenty-four hours before the electrotransfer of DNA, 1 × 10^4^ cells suspended in 500 µl of culture medium were plated in one well of a four-well glass-bottom chambered coverslip (µ-slide, Ibidi) with a growth area per well of 2.5 cm^2^.

### DNA staining and electrotransfer protocol

10 µg of DNA fragments (NoLimits DNA, Thermofisher) at a concentration of 0.5 µg/µl in 10 mM Tris-HCl (pH 7.6) and 1mM EDTA is stained using YOYO-1 dye (1mM in DMSO, Thermofisher) at a bp:YOYO-1 dye molecule staining ratio of 5:1. The stained DNA fragments are then dissolved in the pulsing buffer (10 mM Na_2_HPO_4_*/*KH_2_PO_4_, 1 mM MgCl_2_, 250 mM sucrose, pH 7.0-7.4) at a concentration of 3.33 µg/ml.

The culture media from the chambered coverslips containing the cells is removed just before DNA electrotransfer. The cells are then washed three times with 500 µl of pulsing buffer. 500 µl of the stained DNA solution in the pulsing buffer (3.33 µg/ml) is added to each of the chambered coverslips. 10 electric pulses of 350 V/cm amplitude and 5 ms duration were applied at a frequency of 1 Hz through stainless steel electrodes placed parallel to each other 3 mm apart. The electrode configuration above is expected to provide uniform electric field lines. The electrodes are connected to a pulse generator (BETA tech) to deliver the electric pulses. After the electric pulses are delivered, the electrodes are removed and left to rest in the room temperature (∼22 ^°^C) for 10 minutes to allow resealing of the cell membrane. 500 µl of culture media is then added to the cells (now in the DNA suspension). 500 µl of the resulting solution is removed and the process is repeated three times to dilute the amount of free DNA in the solution. The cells are then placed back in the incubator for 15 minutes.

### DNA tracking experiments

The fluorescence imaging experiments for tracking DNA cargo are performed on an inverted fluorescence microscope (Zeiss Axio-Observer Z1) coupled with an EM-CCD camera (Andor ixon3) with a resolution of 512×512 pixels. The stage of the microscope is coupled to an incubation system (Ibidi) which maintains the cells at 37 ^°^C, 5% CO_2_, and *>* 90 % humidity. A 100×/1.2 oil immersion objective (Zeiss Acroplan) magnifies the image to a field of view of 81.92 × 81.92 µm^2^. An HXP 120C lighting unit (Zeiss, Germany) coupled with a ET-EGFP filter set (excitation: 470 nm/emission: 525 nm) is used as the light source for fluorescence imaging of YOYO-1 stained DNA. The trajectory of internalized DNA is recorded as a time series of 350 images (14 bit) with an exposure time of 100 ms at a camera streaming mode, resulting in a frequency of 10 frames per second. The images were acquired by focusing on a plane with sharp fluorescent DNA cargo. The recorded trajectory is a two dimensional projection of the trajectory of the DNA cargo. The tracking experiments are done between 20 minutes and 1 hour after the application of electric pulses.

### Data analysis

One to five DNA aggregates were observed per cell after electrotransfer. The background noise and larger particles in the image time series is removed by applying an appropriate band pass filter through a home-made script written in Python. The position **r** of the individual DNA cargo are detected using customized version of Trackpy package written in Python [45]. The analysis of the data is then performed by home-made scripts written in Python. The following packages in Python were used in this research: Trackpy, Numpy, Scipy, Pandas and Matplotlib.

### Statistical analysis

The statistical analysis was performed using one way analysis of variance (ANOVA) in Python. The statistical analysis was further confirmed using two sided T tests in Python. *: *P* < 0.05, **:*P* < 0.01, ***: *P* < 0.001, ****: *P* < 0.0001.

## ACKNOWLEDGMENTS

The authors thank Fabio Grillo (ETH Zurich, Switzerland) and Siddhartha Mukherjee (ICTS, Bengaluru, India) for the insightful discussions. The authors also acknowledge Isabell Bagemihl and Reece Lewis (TU Delft, the Netherlands) for the critical reading of the manuscript. The research performed in this article is supported by the funding from the European Research Council (ERC) under the European Union’s Horizon 2020 research and innovation programme (grant agreement no. 819424).

