## Supplementary information for "Intracellular transport of electrotransferred DNA cargo is governed by coexisting ergodic and non ergodic anomalous diffusion"

Aswin Muralidharan,[1,*] Hans Uitenbroek,[1] and Pouyan E. Boukany[1,†]
[1]*Department of Chemical Engineering, Delft University of Technology, van der Maasweg 9, 2629 HZ Delft, the Netherlands*

### Intracellular transport of different sized DNA cargo in CHO-K1 cells

#### Sample trajectories

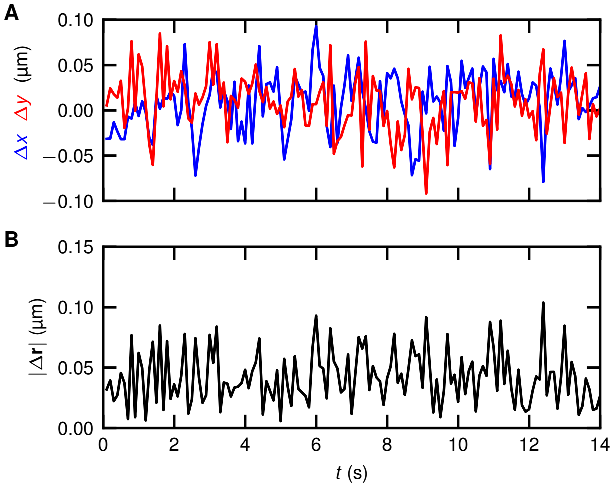


FIG. 1: Representative intracellular DNA cargo displacements in CHO cells. *(A)* Representative $x$ (blue) and $y$ displacements (time interval of 0.1 s) during a measurement. *(B)* Corresponding magnitude of the two dimensional displacement defined as $\Delta\mathbf{r} = |\sqrt{\Delta x^2 + \Delta y^2}|$.

An example of representative intracellular DNA cargo displacement in CHO cells is plotted in Fig. 1. The one dimensional displacements is shown in Fig. 1*A* and the Fig. 1*B* shows the magnitude of the two dimensional displacement defined as $\Delta\mathbf{r} = |\sqrt{\Delta x^2 + \Delta y^2}|$ during a time interval of 0.1 s. We see no evidence of stalling events (zero displacement) that would be seen in a CTRW.

#### Ergodicity breaking

The dependence of the time averaged mean square mean square displacement $\overline{\Delta\mathbf{r}^2(\Delta t, T)}$ with measurement time $T$ is then tested. $\overline{\Delta\mathbf{r}^2(\Delta t, T)}$ was evaluated for different values of $T$ and was plotted in Figs. 4*A*-*C* for different DNA sizes. We extract the power law dependence of $\overline{\Delta\mathbf{r}^2(\Delta t, T)}$ with $T$ by fitting data with the equation $\overline{\Delta\mathbf{r}^2(\Delta t, T)} = AT^{\beta}$, where $\beta$ is the power law exponent and $A$ is a dummy variable. The fitting is performed between $T = 0.8$ s and 10 s to avoid noise at the short time scales due to poor statistics. While the peak of probability density distributions plotted in Fig. 2 is at $\overline{\Delta\mathbf{r}^2(\Delta t, T)} \sim T^0$, non zero power law exponents are present.

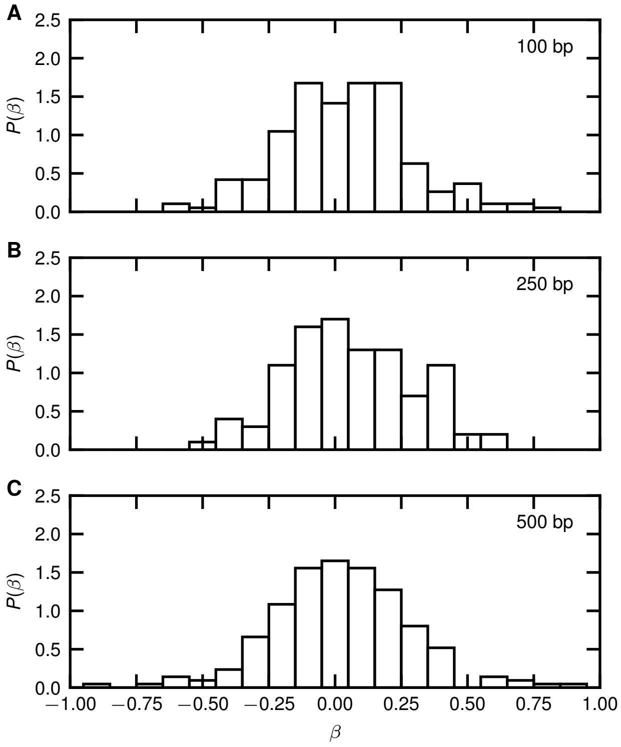


FIG. 2: Probability density distributions of power law exponent $\beta$ obtained by fitting $\overline{\Delta\mathbf{r}^2(\Delta t, T)} \propto T^{\beta}$ is plotted for *(A)* 100 bp, *(B)* 250 bp, *(C)* 500 bp DNA. The fitting is performed between measurement time $T = 0.8$ s and 10 s to avoid errors due to poor statistics at short time scales.

#### Directional change distribution

We plot the directional change probability density distribution between steps for the different sizes of DNA in Figs. 3*A*-*C*. The relative angle $\theta(t, \Delta t)$ is defined as

$$\cos\theta(t, \Delta t) = \frac{\Delta\mathbf{r}(t) \cdot \Delta\mathbf{r}(t + \Delta t)}{|\Delta\mathbf{r}(t)|\,|\Delta\mathbf{r}(t + \Delta t)|}, \qquad (1)$$

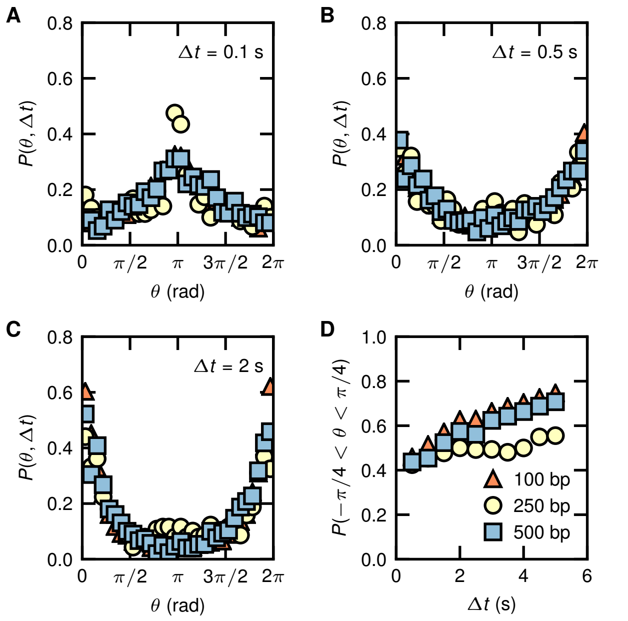


FIG. 3: Directional change distribution for 100 bp, 250 bp and 500 bp DNA for different degrees of temporal coarsening *(A)* $\Delta t = 0.1$ s, *(B)* $\Delta t = 0.5$ s, and *(C)* $\Delta t = 2$ s. *(D)* Probability of increase in directionality ($P(-\pi/4 < \theta < \pi/4)$) for different DNA sizes is plotted as a function of increasing temporal coarsening $\Delta t$.

where the lag time $\Delta t$ represents the temporal coarsening [1]. At short lag times ($\Delta t = 0.1$ s), the directional change has a peak centered around $\pi$ rad indicating that the DNA is trapped within a confining potential [2]. This is expected as the DNA is trapped in the viscoelastic cytoskeletal meshwork. At further temporal coarsening ($\Delta t = 0.5$ s and $\Delta t = 1$ s), the peak shifts towards 0 rad and $2\pi$ rad showing the presence of directional inertial motion as shown in Figs. 3*B*-*C*. The development of the directionality is then quantified by plotting $P(-\pi/4 < \theta < \pi/4)$ as a function of temporal coarsening in Fig. 3*D*. We see that for all the DNA fragment sizes, directionality of the trajectories increases with increasing temporal coarsening.

#### Apparent diffusion coefficient and anomalous exponent

The two dimensional probability density of logarithm of the apparent diffusion coefficient ($D_{\mathrm{app}}$) and anomalous exponent ($\alpha$) obtained from individual trajectories are plotted in Fig. 4. The data is plotted separately for short lag times in Fig. 4*A*-*C* and long lag times in Fig. 4*D*-*F*.

#### Classification of trajectories

We classify the the trajectories as caged ($\alpha \leq 0.4$), subdiffusive ($0.4 < \alpha \leq 1$), and superdiffusive ($\alpha > 1$) based on value of the anomalous exponent. Fig. 5 shows that $\sim 60\%$ of the trajectories remain caged for all DNA sizes at short lag times. Approximately 35% of the trajectories are subdiffusive and rest superdiffusive at short lag times. At longer lag times, $\sim 20\%$ trajectories remain caged, $\sim 60\%$ trajectories subdiffusive and rest superdiffusive.

#### Velocity autocorrelation

We evaluate $C_v^\delta(\Delta t) = \langle \overline{\mathbf{v}(t+\Delta t)\cdot\mathbf{v}(t)} \rangle$ with different values of $\delta$ to identify if the negative correlation arises from localization errors, confinement or medium elasticity[3]. The $C_v^\delta(\Delta t) = \langle \overline{\mathbf{v}(t+\Delta t)\cdot\mathbf{v}(t)} \rangle$ for 100 bp DNA cargo with $\delta = 0.1, 0.2, 0.5$ s is plotted in Fig. 6. We observe that the $C_v^\delta(\Delta t) = \langle \overline{\mathbf{v}(t+\Delta t)\cdot\mathbf{v}(t)} \rangle$ is self similar and collapses to a single curve for different values of $\delta$. This shows the observed negative correlation does not come from the localization errors but ergodic FBM.

#### Intracellular transport of 500 bp DNA cargo in different cancer cells

#### Non ergodicity of 500 bp DNA cargo in different cancer cells

The heterogeneity in $\overline{\Delta \mathbf{r}^2(\Delta t, T)}$ for $T = 10$ s is presented in Fig. 7*A*-*C* for MDA-MB-231, MCF7 and MCF10A cells. To understand the origin of observed heterogeneity, we checked for ergodicity breaking. We observe that $\overline{\Delta \mathbf{r}^2(\Delta t, T)}$ as presented by its power law dependence with $T$ in Fig. 8*A*-*C*.

#### Apparent diffusion coefficient and anomalous exponent

The two dimensional probability density of logarithm of the apparent diffusion coefficient ($D_{\mathrm{app}}$) and anomalous exponent ($\alpha$) obtained from individual trajectories for different cancer cells are plotted in Fig. 9. The data is plotted separately for short lag times in Fig. 9*A*-*C* and long lag times in Fig. 9*D*-*F*.

#### Classification of trajectories

We classify the the trajectories as caged ($\alpha \leq 0.4$), subdiffusive ($0.4 < \alpha \leq 1$), and superdiffusive ($\alpha > 1$) based on value of the anomalous exponent. Fig. 10 shows that

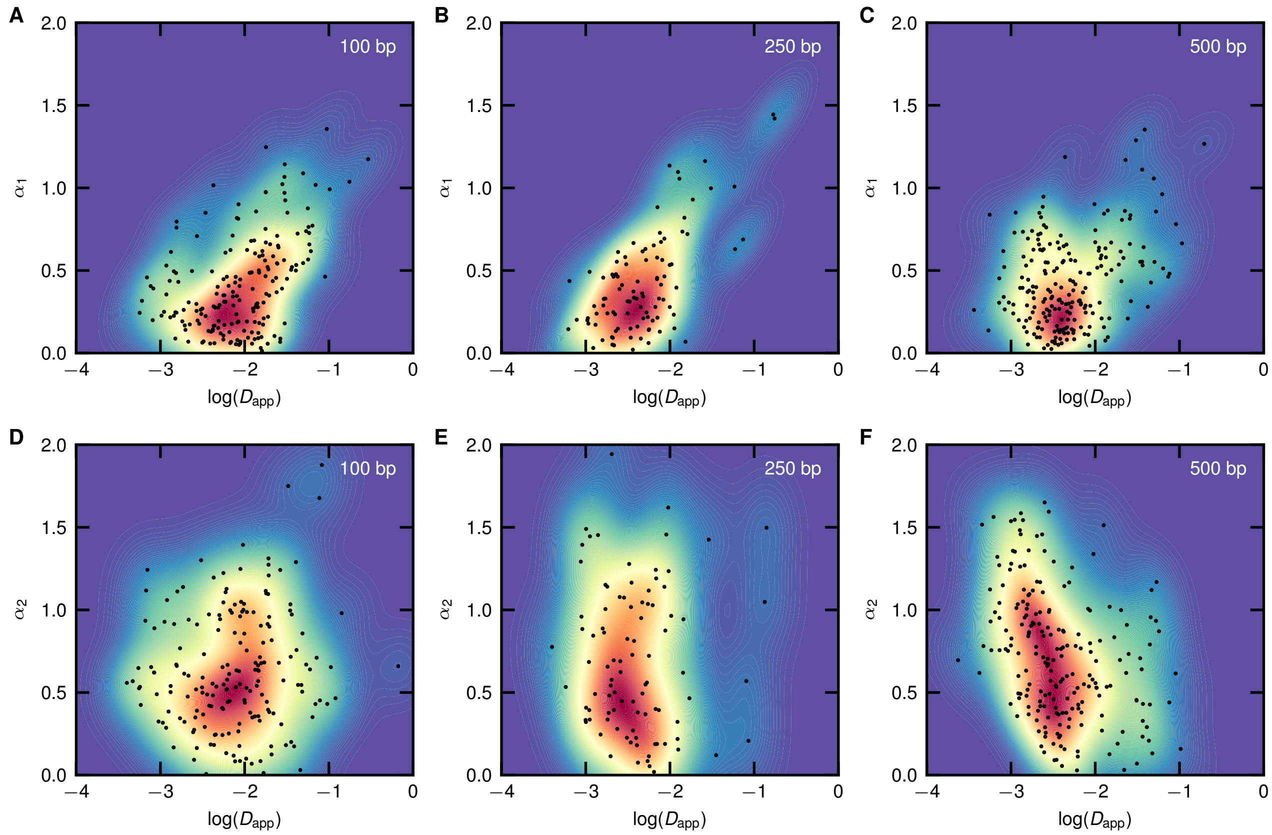


FIG. 4: Scatter plot of logarithm of the apparent diffusion coefficient ($D_{\rm app}$) and anomalous exponent ($\alpha$) obtained at short lag time *(A)-(C)* and long lag time *(D)-(F)* for *(A)*, *(D)* 100 bp, *(B)*, *(E)* 250 bp, and *(C)*, *(F)* 500 bp DNA. The probability density is plotted as a contour with red indicating high density and blue indicating low density.

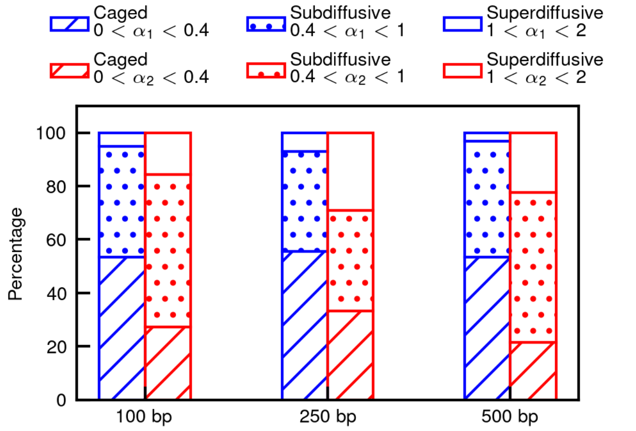


FIG. 5: Classification of individual trajectories to caged ($\alpha \leq 0.4$), subdiffusive ($0.4 < \alpha \leq 1$), and superdiffusive ($\alpha > 1$) for short lag times (blue) and long times (red) for different DNA sizes.

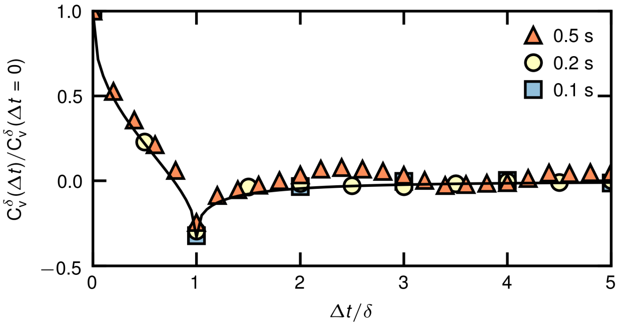


FIG. 6: Velocity auto-correlation function $C_v^\delta(\Delta t) = \langle \mathbf{v}(t+\Delta t)\cdot\mathbf{v}(t)\rangle$, where $\mathbf{v}(t) = [\mathbf{r}(t+\delta) - \mathbf{r}(t)]/\delta$ normalized by $C_v^\delta(\Delta t = 0)$ plotted against the lag time $\Delta t$ normalized by the discretization time interval $\delta = 0.1, 0.2, 9.5$ s for 100 bp DNA cargo in CHO-K1 cell cytoplasm. The solid line is the analytical prediction from fractional Brownian motion (FBM)

$\sim$ 60% of the 500 bp DNA cargo trajectories for MCF7 and MCF10A cells, and $\sim$ 50% of the trajectories and MDA-MB-231 cells remain caged at short lag times. Approximately 35% of the trajectories are subdiffusive and rest superdiffusive at short lag times. At longer lag times, $\sim$ 30% trajectories in MCF10A, $\sim$ 20% trajectories in

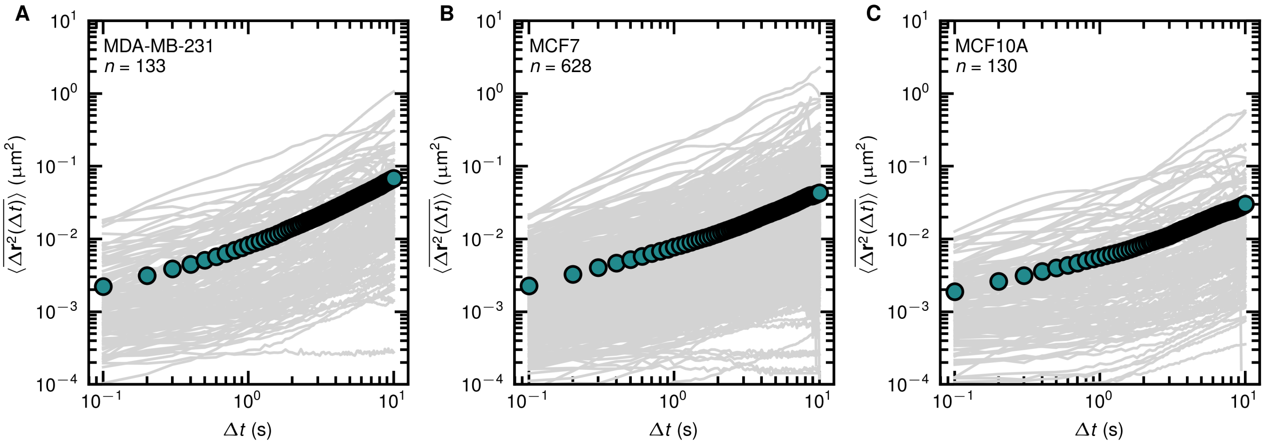


FIG. 7: Individual time averaged mean square displacements ($\overline{\Delta \mathbf{r}^2(\Delta t, T)}$) of 500 bp DNA cargo in cytoplasm are plotted in solid gray color for (*A*) MDA-MB-231 (*B*) MCF7 (*C*) MCF10A cells against the lag time $\Delta t$, for a measurement time $T = 10$ s. The green circles represent the ensemble and time average mean square displacement ($\langle\overline{\Delta \mathbf{r}^2(\Delta t, T)}\rangle$).

MCF7, and $\sim$ 15% trajectories for MDA-MB-231 cells remain caged. $\sim$ 50% of the 500 bp DNA cargo trajectories in MCF10A and MCF7 and $\sim$ 35% in MDA-MB-231 cells are subdiffusive and rest superdiffusive.

#### Directional change distribution

We plot the directional change distribution for 500 bp DNA cargo for MCF10A, MCF7 and MDA-MB-231 cells at different degree temporal coarsening in Fig. 11*A*-*C*. We observe similar behavior in all cell lines that at probability of directionality increases with the increasing temporal coarsening.

* Electronic address: ``
† Electronic address: ``

[1] S. Burov, S. A. Tabei, T. Huynh, M. P. Murrell, L. H. Philipson, S. A. Rice, M. L. Gardel, N. F. Scherer, and A. R. Dinner, Proceedings of the National Academy of Sciences **110**, 19689 (2013).
[2] S. Burov, J.-H. Jeon, R. Metzler, and E. Barkai, Physical Chemistry Chemical Physics **13**, 1800 (2011).
[3] S. C. Weber, M. A. Thompson, W. E. Moerner, A. J. Spakowitz, and J. A. Theriot, Biophysical Journal **102**, 2443 (2012).

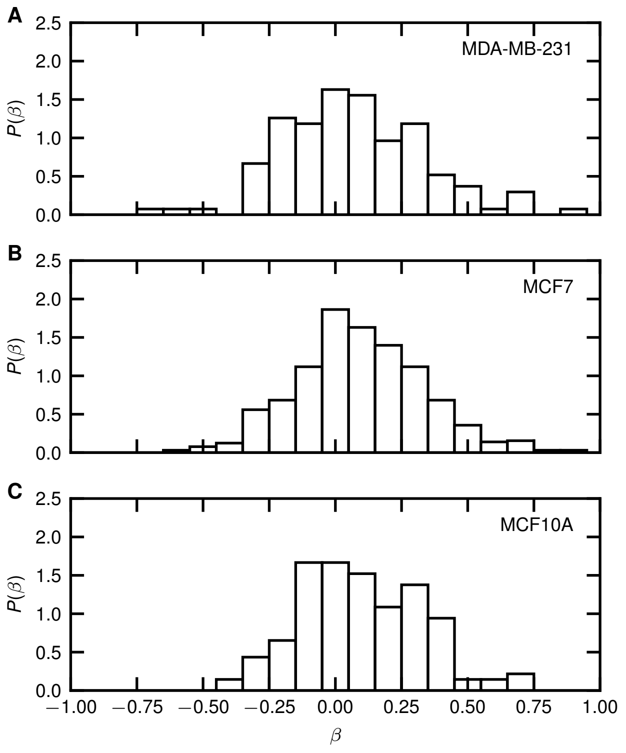


FIG. 8: Probability density distributions of power law exponent $\beta$ obtained by fitting $\overline{\Delta\mathbf{r}^2(\Delta t, T)} \propto T^{\beta}$ of 500 bp DNA cargo in the cytoplasm is plotted for *(A)* MDA-MB-231, *(B)* MCF7, *(C)* MCF10A cells. The fitting is performed between measurement time $T = 0.8$ s and 10 s to avoid errors due to poor statistics at short time scales.

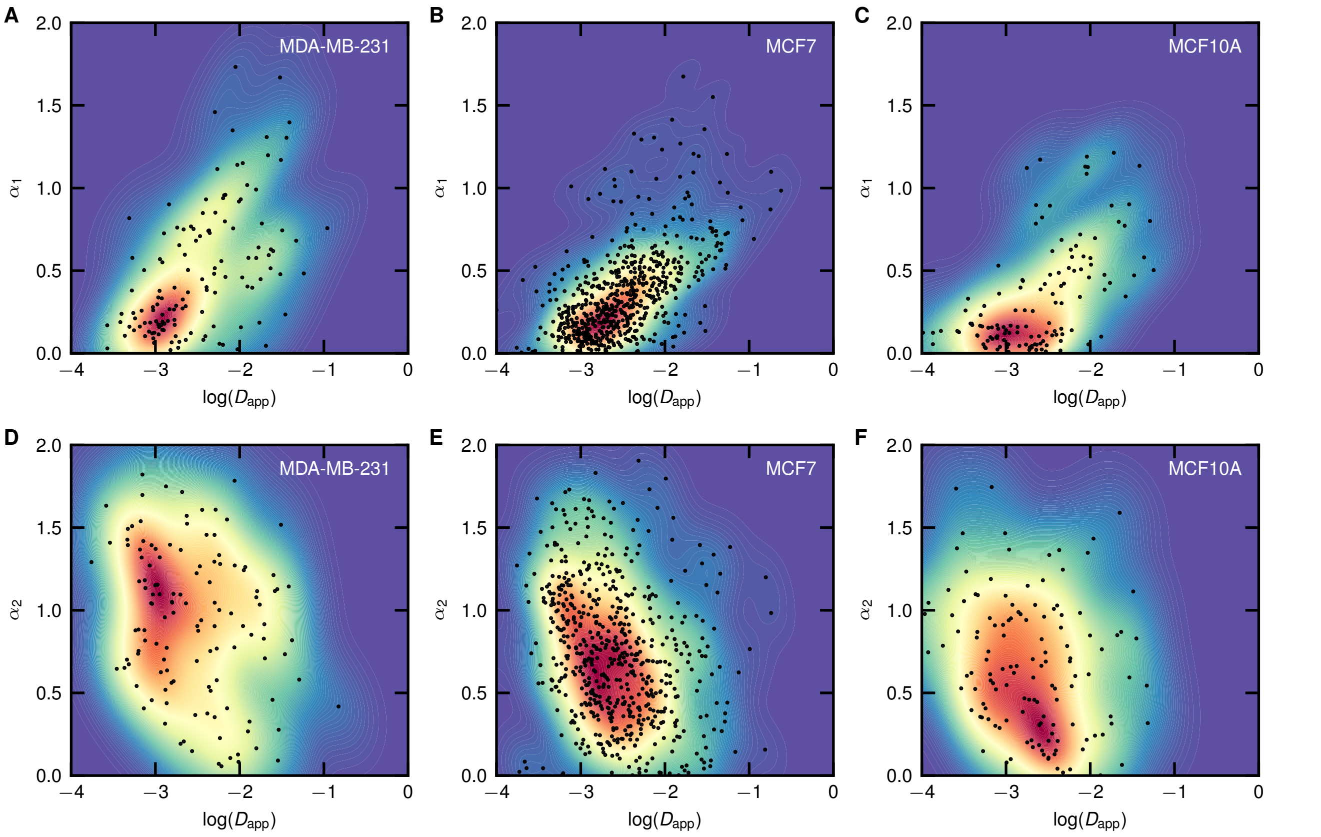


FIG. 9: Scatter plot of logarithm of the apparent diffusion coefficient ($D_{\mathrm{app}}$) and anomalous exponent ($\alpha$) obtained at short lag time *(A)-(C)* and long lag time *(D)*-(F) for *(A)*, *(D)* MDA-MB-231, *(B)*, *(e)* MCF7 bp, and *(C)*, (F) MCF10A cells. The probability density is plotted as a contour with red indicating high density and blue indicating low density.

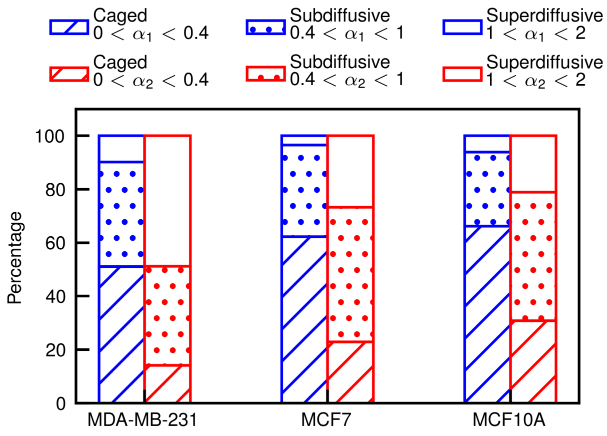


FIG. 10: Classification of individual trajectories to caged ($\alpha \leq 0.4$), subdiffusive ($0.4 < \alpha \leq 1$), and superdiffusive ($\alpha > 1$) for short lag times (blue) and long times (red) for different cancer cells.

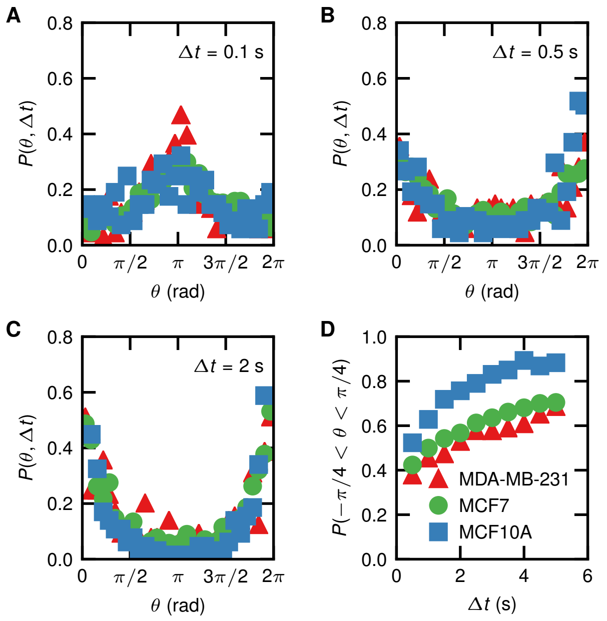


FIG. 11: Directional change distribution of 500 bp DNA cargo in the cytoplasm of MDA-MB-231, MCF7 and MCF10A cells for different degrees of temporal coarsening *(A)* $\Delta t = 0.1$ s, *(B)* $\Delta t = 0.5$ s, and *(C)* $\Delta t = 2$ s. *(D)* Probability of increase in directionality ($P(-\pi/4 < \theta < \pi/4)$) for different DNA sizes is plotted as a function of increasing temporal coarsening $\Delta t$.
